# Fate tracing reveals differences between Reelin^+^ HSCs and Desmin^+^ HSCs in activation, migration, and proliferation activities

**DOI:** 10.1101/2022.02.06.479312

**Authors:** Ning Chen, Shenghui Liu, Dan Qin, Dian Guan, Yaqing Chen, Chenjiao Hou, Songyun Zheng, Liqiang Wang, Xiangmei Chen, Lisheng Zhang

## Abstract

The activation of hepatic stellate cells (HSCs) which comprise distinct clusters, is the main cause of liver fibrogenesis in response to different etiologies of chronic liver injuries. In this study, we constructed a novel ReelinCreERT2 transgenic mouse in which cells expressing Reelin were fully marked and demonstrated that about 50% HSCs were labeled. These ReelinCreERT2-labeled HSCs displayed distinct characteristics in migration, activation and proliferation compared to Desmin^+^ HSCs (total HSCs) in cholestatic (bile duct ligation; BDL) or hepatotoxic (carbon tetrachloride; CCl_4_) liver injuries. In BDL-induced fibrotic livers, Desmin^+^ HSCs were activated with increased proliferation and accumulation activities around the portal triad, but mGFP^+^ HSCs did not show proliferation or accumulation activity around the portal triad, and only a small part was activated. In CCl_4_-induced fibrotic livers, most of Desmin^+^ and mGFP^+^ HSCs were activated along with proliferation and accumulation potential around the central vein, however fewer mGFP^+^ HSCs were activated compared to Desmin^+^ HSCs. Moreover, in the regression of CCl_4_-induced fibrosis, mGFP^+^ HSCs were apoptosed whereas Desmin^+^ HSCs recovered to normal state. Besides, we didn’t find evidence that mGFP^+^ HSCs transdifferentiated into hepatocytes or cholangiocytes through mesenchymal-epithelial transition (MET).

## Introduction

Reelin, a secreted extracellular glycoprotein with a molecular weight of 420 kDa, owns two cell surface receptors, the very low-density lipoprotein receptor and the apolipoprotein E receptor (1–3). In vivo and vitro studies show that Reelin is expressed in hepatocytes and up-regulated in patients with liver cirrhosis in the liver and plasma(2, 4). However, some studies reveal that Reelin is expressed both in hepatocytes and hepatic stellate cells (HSCs)(5, 6). Others demonstrate that Reelin is expressed only in HSCs, but not in hepatocytes(7, 8). Besides these, studies show that Reelin is detected in hepatoblasts and oval cells, which differentiate into hepatocytes and cholangiocytes following liver damage(9). Thus, there exists a huge debate over Reelin localization in liver cells and few detailed functions of Reelin in livers have been investigated.

HSCs belonging to mesenchymal cells exhibit fibroblast and pericyte characteristics, and compose one third of nonparenchymal cells and 15% of resident cells(10, 11). In normal livers, HSCs laden with retinoid droplets maintain quiescent state and are located in the sinusoidal space of Disse where hepatocytes exchange biomolecules with portal blood(12, 13). Following liver injury, quiescent HSCs are activated and transdifferentiated into migratory, contractile, and proliferative myofibroblasts (MFs) to secrete extracellular matrix (ECM)(14, 15). HSC activation leads to fiber scars accumulation in the space of Disse, which further results in endothelial fenestration loss(12, 16). Numerous specific markers, such as desmin, lecithin-retinol acyltransferase (LRAT), collagen type I (Col1a1), vimentin, glial fibrillary acidic protein (GFAP), smooth muscle actin (α-SMA) and cytoglobin have been used to characterize HSCs genetic targeting, imaging, histological detection, and cell fate tracing(17). Genetic cell lineage tracking labeling of MFs with Col1a1, a major component of the extracellular matrix, has showed that HSCs are the primary source of MFs, occupying approximately 87% of MFs in carbon tetrachloride (CCl_4_)-induced liver injury(18). Another study has shown that 82% ~ 96% of MFs are originated from HSCs labeled with LRAT in mice treated with CCl_4_, 3,5 diethoxycarbonyl-1,4-dihydro-collidin diet or bile duct ligation (BDL)(19). All of these suggest that HSCs are the major source of MFs and the activation of HSCs is the main cause of liver fibrogenesis in response to diverse etiologies.

Several studies have reported that mesenchymal-to-epithelial transition (MET) occurs following liver injury and chronic liver inflammation to reduce fibrosis(20–22). Studies using GFAPCre and ACTA2CreERT2-marked mice have demonstrated that HSCs differentiate into hepatocytes and cholangiocytes through MET in injured livers(23–25). Other studies using LratCre- and VimentinCreER-labeled mice have documented no HSCs undergo MET during liver injury(19, 26, 27). So far, whether HSCs undergo MET is still a scientific question to be addressed. After underlying etiology of liver fibrosis is removed, fibrosis scars are gradually regressed(10, 13, 15). In the course of this regression, the activated HSCs undergo apoptosis or revert to a quiescent-like state(28, 29). The apoptosed and inactivated HSCs in the regression of liver fibrosis may be originate from different subsets(30). Promoting HSC apoptosis and repressing HSC activation through pharmacological treatment contributes to liver fibrosis resolution (31). Therefore, facilitating HSC apoptosis and repressing HSC activation have been viewed as a therapeutic target for liver fibrosis(32, 33).

Although HSCs play a major role in response to various types of liver fibrosis, the fibrogenic phenotype and mechanisms are different according to the various kinds of etiologies(34, 35). Therefore, a variety of models, such as BDL and CCl_4_, are used to mimic hepatopathy of different types to develop better therapeutic strategies (36). BDL and CCl_4_ induced liver injuries are two distinct liver fibrosis models that mimics cholestasis (such as primary sclerosing cholangitis and primary biliary cirrhosis) and hepatotoxicity (such as nonalcoholic steatohepatitis and chronic viral hepatitis) respectively(37, 38). Additionally, single-cell RNA sequencing reveals that HSCs are heterogeneous and comprised of distinct populations with different gene-expression in normal and a variety of disease livers and divided into different clusters according to their distinct characteristics of position and function(30, 39, 40). These findings indicate that liver fibrosis treatment should suit the remedy to the case based on fibrogenetic etiologies in different etiologies of liver disease(14).

In this study, our genetic cell fate tracking data revealed that ReelinCreERT2-labeled HSCs displayed different characteristics compared to Desmin^+^ HSCs (total HSCs) in BDL-induced and CCl_4_-induced fibrotic livers.

## Results

### ReelinCreERT2 labels HSCs in sham-operated and BDL-induced fibrotic mouse livers

In order to achieve accurate labelling of Reelin-expressing cells, we constructed Reelin^CreERT2^; Rosa26mTmG^flox^ (R26T/G^f^) mouse model. In this model, after TAM treatment, tomato sequence was excised by Cre, and membrane-tagged green fluorescence protein (mGFP) started to be expressed (**Figure 1A**). When Reelin^CreERT2^; R26T/G^f^ mice were injected with TAM, ReelinCreERT2-marked cells in the livers expressed mGFP (**Figure 1B**). Then, we verified whether ReelinCreERT2-mediated mGFP expression matched endogenous Reelin expression in sham-operated and BDL-induced fibrotic mouse livers. Obvious collagen fiber was accumulated after BDL operation (**Supplemental Figure 1A**), and α-SMA and Col1a1 expression was significantly increased (**Supplemental Figure 1B**) compared to those in sham-operated livers, indicating livers developed significant fibrosis following BDL operation. Immunostaining of mGFP and Reelin in serial section indicated that ReelinCreERT2-mediated mGFP expression almost fully matched endogenous Reelin both in sham-operated and BDL-induced fibrotic mouse livers (**Figure 1C**). These findings suggest that Reelin^CreERT2^; R26T/G^f^ mouse was a credible model to precisely label cells expressing Reelin. Next, we explored what types of cells Reelin located in in mouse livers. Immunohistochemistry of the adult Reelin^CreERT2^; R26T/G^f^ mouse livers for mGFP and Desmin showed almost all of mGFP^+^ cells were Desmin^+^ but some Desmin^+^ cells were not mGFP^+^ (**Figure 1D**), indicating that Reelin was expressed in HSCs and only part of HSCs expressed Reelin rather than hepatocytes, hepatoblasts or oval cells in sham-operated and BDL-induced fibrotic mouse livers.

**Figure 1.**
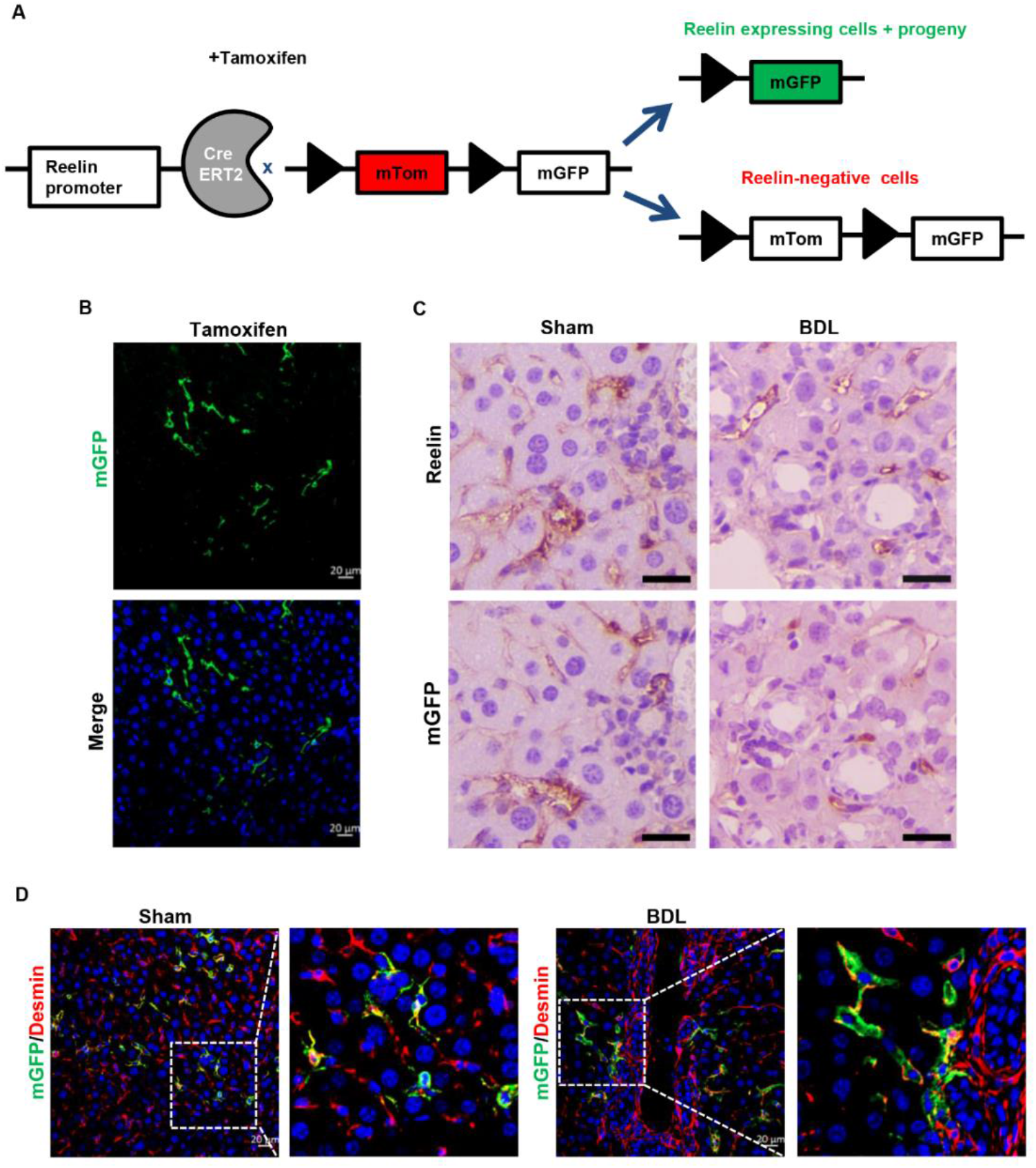
Reelin is expressed in HSCs in the sham and BDL treated livers. **A.** Schematic diagram showing mTom/mGFP reporter gene expression in the absence and presence of tamoxifen-inducible CreERT2-mediated recombination. **B.** After treated with TAM, mGFP was induced in Reelin^CreERT2^; R26T/G^f^ mice. **C.** Livers from sham or BDL-operated Reelin^CreERT2^; R26T/G^f^ mice were stained with anti-Reelin and anti-GFP antibodies, and analyzed immunohistochemistry demonstrating overlap between ReelinCreERT2-induced mGFP expression and endogenous Reelin expression in serial sections. **D.** Immunohistochemistry of the sham or BDL-operated adult Reelin^CreERT2^; R26T/G^f^ mouse livers for mGFP with Desmin demonstrated that Reelin was expressed in HSCs and only part of HSCs expressed mGFP. Scale bar in B and D represents 20 μm. Scale bar in C represents 50 μm.

### ReelinCreERT2-labeled HSCs do not migrate or accumulate and only a small fraction is activated in BDL-induced fibrotic livers

As Reelin was expressed only in part of HSCs by immunostaining of mGFP and Desmin, we investigated whether there were differences between Desmin^+^ HSCs and mGFP^+^ HSCs. Immunohistochemistry of mGFP and Desmin with Glutamine Synthetase (GS, a marker of central vein) showed that both mGFP^+^ and Desmin^+^ HSCs were scattered throughout the parenchyma in sham-operated livers (**Figure 2A**). Whereas in BDL-induced fibrotic livers, Desmin^+^ HSCs were accumulated around the portal triad, and mGFP^+^ HSCs were scattered throughout the parenchyma (**Figure 2B**). These findings suggest significant differences exist in migration capacity and location between Desmin^+^ and mGFP^+^ HSCs in BDL-induced liver fibrosis. Next, we explored activation ability of mGFP^+^ HSCs. Immunostaining of mGFP with α-SMA showed that mGFP^+^ HSCs did not express α-SMA in sham-operated livers, however, α-SMA was expressed in mGFP^+^ HSCs in BDL-induced fibrotic livers (**Figure 2C**), which indicated that mGFP^+^ HSCs were activated in fibrotic livers induced by BDL. Immunostaining of α-SMA with Desmin and mGFP in serial sections showed that a large proportion of Desmin^+^ HSCs (72.35% Desmin^+^ HSCs) expressed α-SMA, but only 31.01% mGFP^+^ HSCs expressed α-SMA in BDL-induced fibrotic livers (**Figure 2D**). To confirm the finding that fewer mGFP^+^ HSCs were activated than Desmin^+^ HSCs, we analyzed serial sections immunostaining of Col1a1 with mGFP or Desmin, and got similar results as shown in **Figure 2E** that Col1a1 was expressed in 48.53% mGFP^+^ HSCs while expressed in 89.34% Desmin^+^ HSCs.

**Figure 2.**
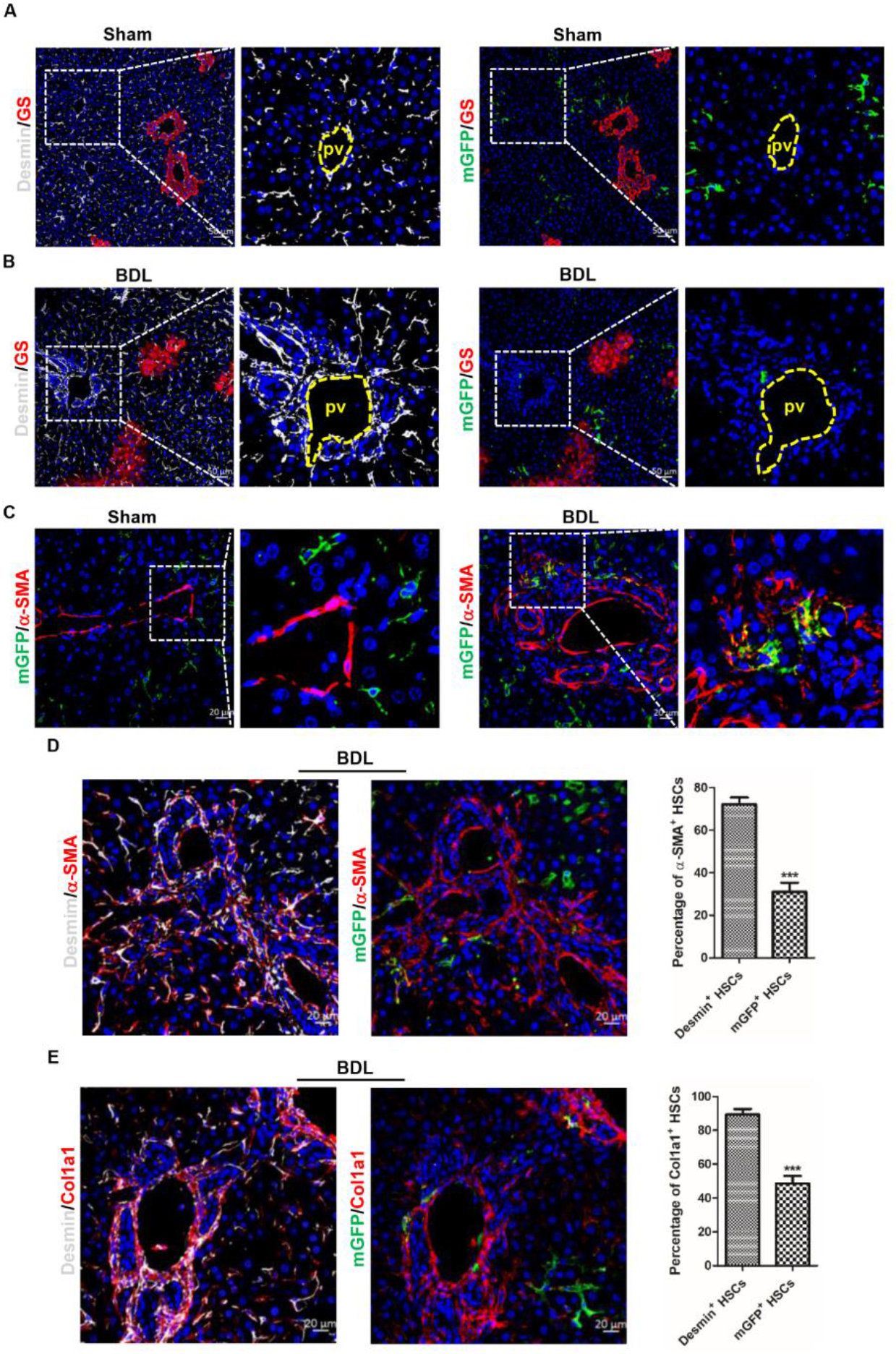
ReelinCreERT2 labeled HSCs do not accumulate and fewer cells are activated compared to Desmin^+^ HSCs in BDL-induced fibrotic livers. **A.** mGFP and Desmin costaining with GS in sham-operated Reelin^CreERT2^; R26T/G^f^ mouse liver determined that mGFP^+^ HSCs and Desmin^+^ HSCs were scattered throughout the parenchyma. **B.** mGFP and Desmin costaining with GS in BDL-operated Reelin^CreERT2^; R26T/G^f^ mouse liver for 2 weeks indicated that Desmin^+^ HSCs accumulated around the portal vein (pv), whereas mGFP^+^ HSCs were scattered throughout the parenchyma. **C.** mGFP^+^ HSCs were activated in BDL-induced Reelin^CreERT2^; R26T/G^f^ mouse fibrotic livers observed by immunohistochemistry of mGFP and α-SMA. **D.** Immunohistochemistry of the BDL-induced fibrotic livers for Desmin with α-SMA and mGFP with α-SMA observed that fewer mGFP^+^ HSCs expressed α-SMA. **E.** Analyzed immunohistochemistry of the BDL-induced fibrotic livers for Desmin with Col1a1 and mGFP with Col1a1 determined that fewer mGFP^+^ HSCs expressed Col1a1. Data are reported as means ± SEM. *p < 0.05; **p < 0.01; ***p < 0.001. Scale bar in A and B represents 50 μm. Scale bar in C, D, and E represents 20 μm.

### ReelinCreERT2-labeled HSCs do not show distinguished proliferation activity in BDL-induced fibrotic livers

The migration, proliferation and activation of HSCs are an accompanying process(10, 12). There were significant differences in migration and activation activities between mGFP^+^ and Desmin^+^ HSCs, so we want to know the proliferative capacity difference between mGFP^+^ and Desmin^+^ HSCs. Immunohistochemical staining of Desmin and mGFP showed the number of Desmin^+^ HSCs increased remarkably in BDL-induced fibrotic livers compared to sham-operated livers (**Figure 3A**), but the number of mGFP^+^ HSCs were comparable (**Figure 3B**). Co-staining of mGFP and Desmin showed the percentage of mGFP^+^ HSCs accounted for Desmin^+^ HSCs was 49.83% in sham-operated livers but decreased to 23.84% in BDL-induced fibrotic livers (**Figure 3C**), which further confirmed that the number of Desmin^+^ HSCs increased remarkably but the number of mGFP^+^ HSCs had no significant difference in BDL-induced fibrotic livers. Bromodeoxyuridine (BrdU) labeling showed the percentage of Desmin^+^ HSCs with BrdU was 4.74% in sham-operated livers and increased to 9.58% in BDL-induced livers, but the percentage of mGFP^+^ HSCs labeled by BrdU did not change much (4.34% in sham-operated livers and 3.31% in BDL-induced livers) (**Figure 3D**). Immunostaining of Ki67 and mGFP or Desmin got a similar result that the proliferation ratio of Desmin^+^ HSCs increased greatly (6.57% in sham-operated livers and increased to 10.59% in BDL-induced fibrotic livers), but the ratio of mGFP^+^ HSCs had no notable difference (5.84% in sham-operated livers and 5.33% in BDL-induced fibrotic livers) (**Figure 3E**). Considering the factors, such as changes of cell number, proliferation rate, and the percentage of mGFP^+^ HSCs accounted for Desmin^+^ HSCs in normal and BDL-induced fibrotic livers, we conclude that the number of Desmin^+^ HSCs increased greatly, but the proliferative ratio of mGFP^+^ HSCs in BDL-induced fibrotic livers was not remarkably different from sham-operated livers.

**Figure 3.**
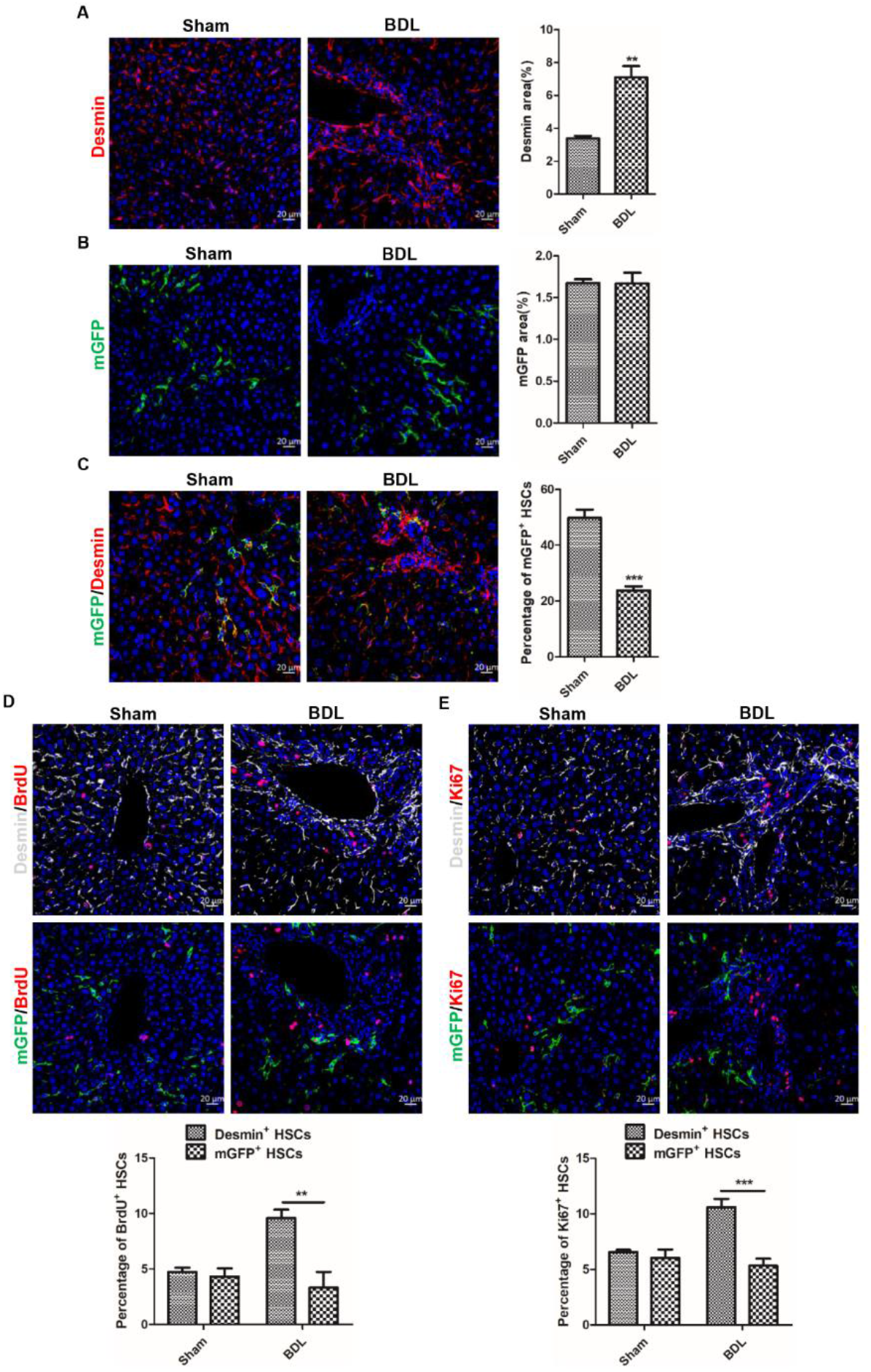
ReelinCreERT2 labeled HSCs do not show proliferation capacity in BDL-induced fibrotic livers. **A.** The number of Desmin^+^ HSCs increased significantly in BDL-induced liver injury observed by immunohistochemistry of Desmin^+^ HSCs. **B.** Analyzed immunohistochemistry of mGFP^+^ HSCs indicated that the number of mGFP^+^ HSCs was not increased in injured livers induced by BDL. **C.** The percentage of mGFP^+^ HSCs accounted for Desmin^+^ HSCs was significantly reduced in Reelin^CreERT2^; R26T/G^f^ mouse fibrotic livers determined by BDL Immunostaining of mGFP and Desmin. **D.** Proliferative properties of Reelin^+^ HSCs and Desmin^+^ HSCs determined by immunohistochemistry for BrdU with Desmin or mGFP demonstrated that Desmin^+^ HSCs proliferative properties significantly increased in BDL-induced liver fibrosis comparing to sham-operated livers, but mGFP^+^ HSCs proliferative properties had no difference. **E.** Immunohistochemistry for Ki67 with Desmin or mGFP demonstrated that Desmin^+^ HSCs proliferative properties significantly increased in BDL-induced liver fibrosis comparing to sham-operated livers, but mGFP^+^ HSCs proliferative properties had no difference. Data are reported as means ±SEM. *p < 0.05; **p < 0.01; ***p < 0.001. Scale bar represents 20 μm.

### mGFP^+^ HSCs accumulate around central vein in CCl_4_-induced liver injury

BDL initially induced biliary duct hyperplasia and further caused biliary fibrosis, however, CCl_4_-induced liver fibrosis started on pericentral cell injury and formed fibrous septum (**Supplemental Figure 1C**). For this reason, we explored whether there were differences in migration, activation and proliferation potentials between mGFP^+^ HSCs and Desmin^+^ HSCs in CCl_4_-induced liver injury. After treated with CCl_4_ for 6 weeks, obvious collagen fiber was observed by Sirius red staining (**Supplemental Figure 2A**) and immunostaining showed that the expression of α-SMA and Col1a1 was significantly increased. (**Supplemental Figure 2B**), which indicated that Reelin^CreERT2^; R26T/G^f^ mice developed severe fibrosis. Serial section immunohistochemistry of Reelin and mGFP in normal and CCl_4_-induced fibrotic livers of TAM-treated Reelin^CreERT2^; R26T/G^f^ mice showed mGFP expression almost fully matched endogenous Reelin, which was similar to immunohistochemistry staining of Reelin and mGFP in sham-operated and BDL-induced fibrotic livers (**Figure 4A**). Immunohistochemistry for mGFP with Desmin and α-SMA also showed mGFP was expressed in HSCs in normal and CCl_4_-induced fibrotic livers (**Figure 4B**) and mGFP^+^ HSCs were activated in CCl_4_-induced fibrotic livers (**Figure 4C**). Besides, we observed that mGFP^+^ and Desmin^+^ HSCs were scattered throughout the parenchyma in normal livers (**Figure 4D**), but both of them were accumulated around the central vein in CCl_4_-induced fibrotic livers (**Figure 4E**). These findings indicate that the migration activity of Reelin^+^ HSCs in CCl_4_-induced fibrotic livers was significantly different with that in BDL-induced fibrotic livers.

**Figure 4.**
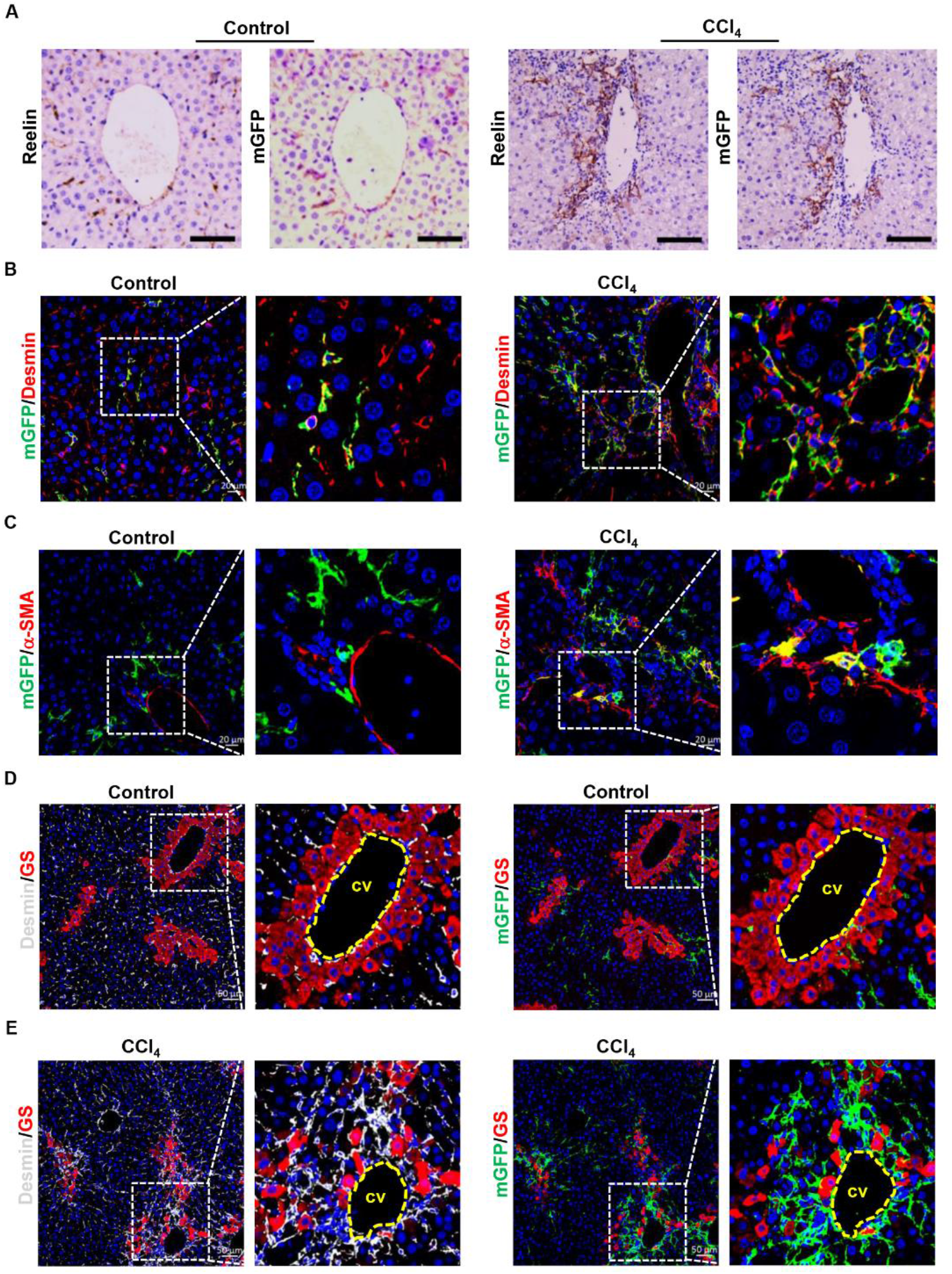
Genetically labeled Reelin^+^ HSCs accumulate around the central veins and fibrous septa in CCl_4_-induced injured livers. **A.** Immunostaining of Reelin and mGFP in vehicle-treated or CCl_4_-treated Reelin^CreERT2^; R26T/G^f^ mouse livers for 6-week demonstrated overlap between endogenous Reelin expression and ReelinCreERT2-induced mGFP expression in serial sections. **B.** Immunohistochemistry of normal or fibrotic livers induced by CCl_4_ for mGFP and Desmin showed that mGFP was located in part of HSCs. **C.** Activated mGFP^+^ HSCs were observed in CCl_4_-induced fibrotic livers by immunostaining of mGFP and α-SMA. **D.** Immunohistochemistry of normal livers for GS with mGFP or Desmin showed mGFP^+^ HSCs and Desmin^+^ HSCs were scattered throughout the parenchyma. **E.** Immunohistochemistry of CCl_4_-induced fibrotic livers for GS with mGFP or Desmin showed both mGFP^+^ HSCs and Desmin^+^ HSCs accumulated around the central veins (cv) and fibrous septa. Scale bar in A represents 100 μm. Scale bar in B and C represents 100 μm. Scale bar in A represents 50 μm.

### mGFP^+^ HSCs share similarity with Desmin^+^ HSCs in proliferation but fewer cells are activated compared to Desmin^+^ HSCs in CCl_4_-induced fibrotic livers

Next, we explored mGFP^+^ HSC’s activation and proliferation properties in CCl_4_-induced fibrotic livers. In CCl_4_-induced fibrotic livers of ReelinCreERT2; R26T/G^f^ mice, the Immunohistochemistry results showed that 60.43% mGFP^+^ HSCs expressed α-SMA and the proportion of Desmin^+^ HSCs expressed α-SMA was 80.37% (**Figure 5A**). Meanwhile, 75.38% mGFP^+^ HSCs and 85.42% Desmin^+^ HSCs expressed Col1a1 (**Figure 5B**). The above results indicated that fewer mGFP^+^ HSCs were activated than Desmin^+^ HSCs in CCl_4_-induced fibrotic livers. And immunostaining of Desmin and mGFP showed that the number of Desmin^+^ HSCs and mGFP^+^ HSCs was increased greatly in CCl_4_-treated fibrotic livers (**Figure 5C and D**). However, co-staining of mGFP and Desmin revealed that the percentage of mGFP^+^ HSCs accounted for Desmin^+^ HSCs had no significant difference in CCl_4_-treated fibrotic livers compared to that in normal livers (**Figure 5E**), which indicated there was no significant difference between Desmin^+^ and mGFP^+^ HSCs in regarding to proliferative property. So, we further investigated the proliferative property of mGFP^+^ HSCs and Desmin^+^ HSCs by BrdU labeling and Ki67 staining in normal and CCl_4_-treated livers. Our results documented that Desmin^+^ and mGFP^+^ HSCs had superior proliferation ability in CCl_4_-treated livers compared to normal livers, but the proliferation rate was comparable between Desmin^+^ and mGFP^+^ HSCs (**Figure 5F and G**). Based on the immunohistochemistry staining for mGFP and Desmin with α-SMA, Col1a1, BrdU and Ki67, we conclude that, in CCl_4_-induced injured livers, mGFP^+^ HSCs shared similarities with Desmin^+^ HSCs in proliferation property but still fewer mGFP^+^ HSCs were activated compared to Desmin^+^ HSCs.

**Figure 5.**
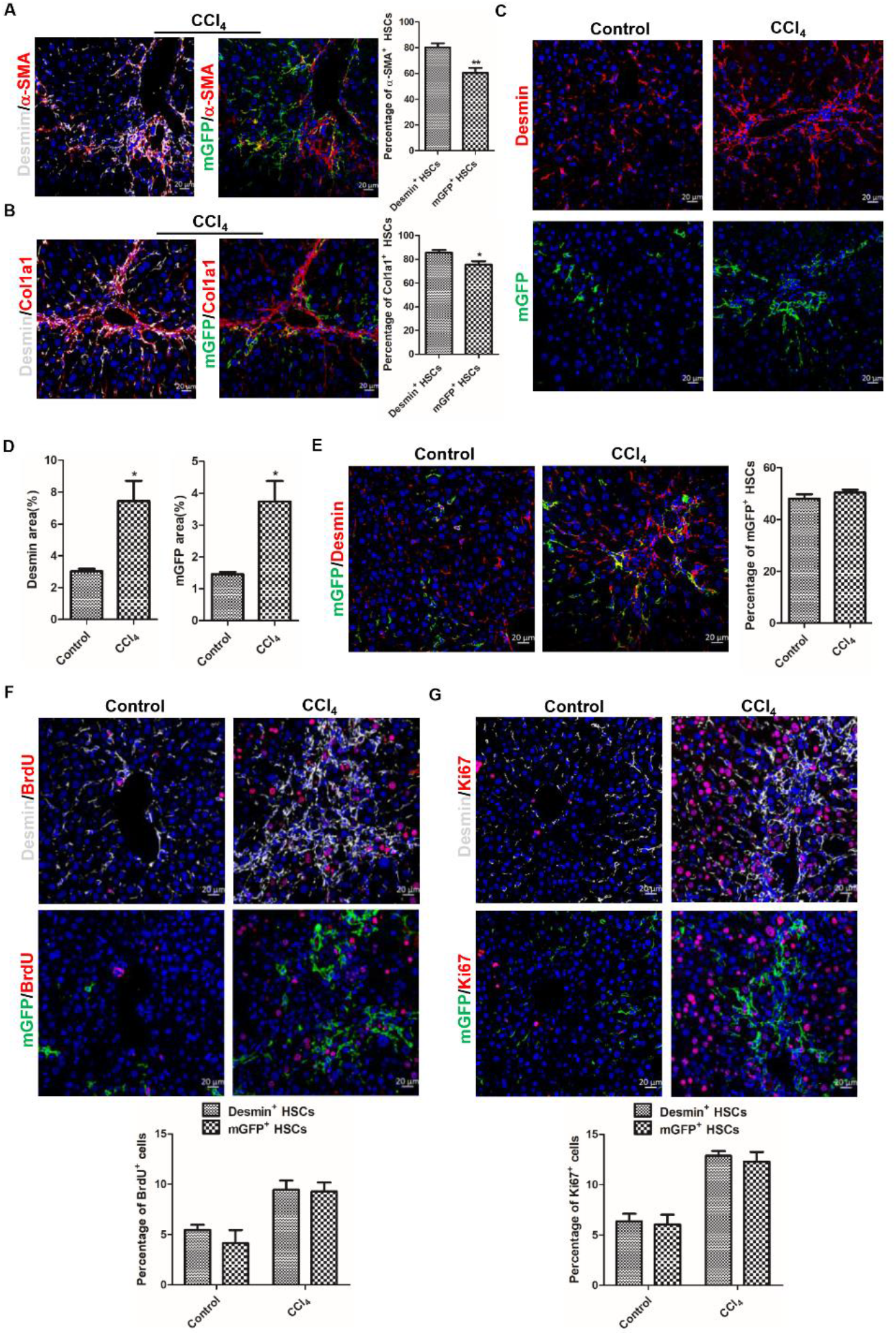
Fewer GFP^+^ HSCs are activated compared to Desmin^+^ HSCs in CCl_4_-induced injured livers. **A.** Analyzed immunohistochemistry of Reelin^CreERT2^; R26T/G^f^ mice treated with CCl_4_ for 6-week for α-SMA with mGFP or Desmin demonstrated that fewer mGFP^+^ HSCs expressed α-SMA. **B.** Immunostaining for Col1a1 with mGFP or Desmin observed that fewer mGFP^+^ HSCs expressed Col1a1. **C**. Immunostaining of mGFP and Desmin displayed that the number of Desmin^+^ HSCs and mGFP^+^ HSCs was significantly increased in CCl_4_-induced fibrotic livers. **D.** Quantification data of Desmin^+^ HSCs and mGFP^+^ HSCs. **E.** Analyzed immunostaining of mGFP and Desmin indicated that the percentage of mGFP^+^ HSCs accounted for Desmin^+^ HSCs had no significant difference in fibrotic livers induced by CCl_4_ compared to in normal livers. **F.** Proliferative properties of Reelin^+^ HSCs were determined by BrdU costaining with desmin or mGFP in serial sections of the mice treated with vehicle or CCl_4_ for 6-week, and Desmin^+^ HSCs and mGFP^+^ HSCs showed significant proliferation activity following CCl_4_ injection. **G.** Proliferative properties of Desmin^+^ HSCs and GFP^+^ HSCs were determined by Ki67 in Desmin^+^ HSCs and GFP^+^ HSCs. Data are reported as means ± SEM. *p < 0.05; **p< 0.01; ***p < 0.001. Scale bar represents 20 μm.

### ReelinCreERT2-labeled HSCs do not transdifferentiate into hepatocytes or cholangiocytes in healthy or injured livers

HSCs transdifferentiating into hepatocytes or cholangiocytes in injured livers is controversial(21, 27, 41). To verify whether HSCs are able to transdifferentiate into hepatocytes and cholangiocytes, we tested ReelinCreERT2-labeled HSCs transformation in sham-operated and BDL-induced fibrotic livers. However, analyzed immunohistochemistry for mGFP with hepatocyte nuclear factor 4 alpha (HNF4α) showed no mGFP^+^ HSCs were observed with typical hepatocyte nuclear morphology or expressed HNF4α both in sham-operated and BDL-induced fibrotic livers (**Figure 6A**). Moreover, immunostaining of mGFP with cytokeratin 19 (CK19) observed no mGFP^+^ HSCs expressed CK19 either (**Figure6B**). Although we did not observe ReelinCreERT2-labeled HSCs transdifferentiated into hepatocytes or cholangiocytes in BDL-induced fibrotic livers, in consideration of the different mechanism between BDL-induced and CCl_4_-induced liver fibrosis, we tested whether ReelinCreERT2-labeled HSCs transdifferentiate into hepatocytes or cholangiocytes through MET in CCl_4_-induced fibrotic livers. Immunostaining of mGFP with HNF4α or CK19 in normal, CCl_4_-treated, CCl_4_-treated following 7 days recovery or CCl_4_-treated following 3 weeks recovery livers showed no mGFP^+^ HSCs expressed HNF4α or CK19 (**Figure 6C and 6D**). Collectively, these findings exclude the possibility that ReelinCreERT2-marked HSCs transdifferentiated into hepatocytes or cholangiocytes through MET in healthy or injured livers.

**Figure 6.**
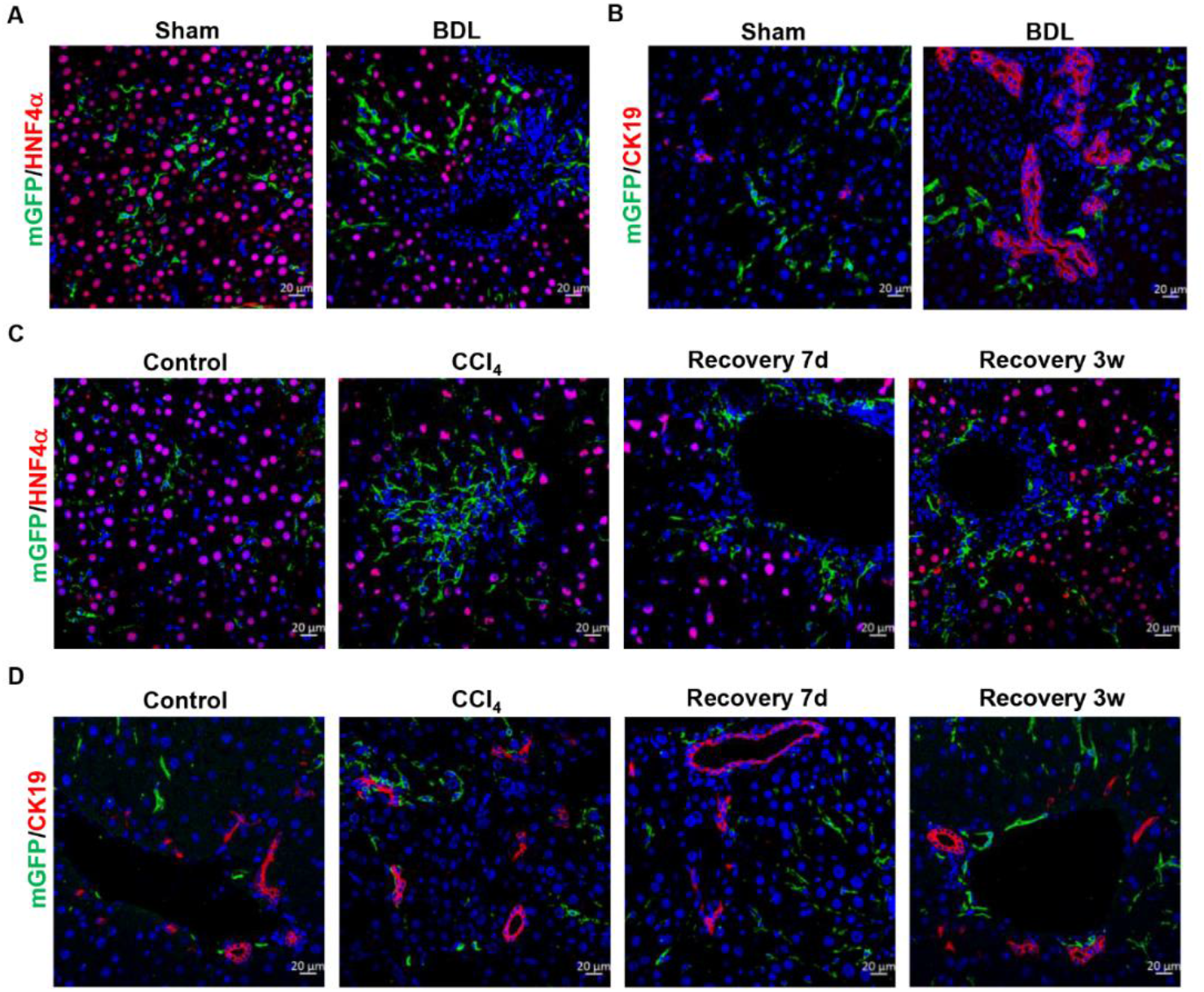
Genetically labeled Reelin^+^ HSCs do not undergo MET in response to chronic liver injury or recovery from fibrosis. **A.** No genetically labeled mGFP^+^ HSCs was observed expressing hepatocytes marker HNF4α in sham-operated livers or BDL-induced fibrotic livers. **B**. Genetically labeled mGFP^+^ HSCs did not express cholangiocytes marker CK19 in sham-operated livers or BDL-induced fibrotic livers. **C.** Genetically labeled mGFP^+^ HSCs in livers treated with vehicle or in livers in response to CCl_4_ for 6-week, recovery from CCl_4_ for 7-day or 3-week did not express hepatocytes marker HNF4α in ReelinCreERT2 mice. **D**. Genetically labeled mGFP^+^ HSCs in normal livers or in livers in response to CCl_4_ for 6-week, recovery 7-day or 3week from CCl_4_ in ReelinCreERT2 mice did not express cholangiocytes marker CK19. Scale bar represents 20 μm.

### ReelinCreERT2-labeled HSCs undergo apoptosis in CCl_4_-induced liver fibrosis regression

We further investigated mGFP^+^ HSC’s cell fate in livers recovered from CCl_4_-induced injury. Sirius red staining showed collagen fiber deposited obviously in CCl_4_-treated livers and regressed markedly in livers recovered from CCl_4_-induced injury (**Supplemental Figure 2C**). Immunostaining of Desmin and mGFP showed, in CCl_4_-induced fibrotic livers, Desmin^+^ and mGFP^+^ HSCs proliferated notably and after 5 week’s recovery from CCl_4_-induced injury the number of Desmin^+^ HSCs returned to normal, whereas few mGFP^+^ HSCs were left (**Figure 7A and B**). The percentage of mGFP^+^ HSCs accounted for Desmin^+^ HSCs was 48.08% in normal livers and 50.49% in CCl_4_-induced fibrotic livers, but decreased to 5.60% in livers recovered from CCl_4_-induced injury for 5 weeks (**Figure 7C**). Early study reported that HSCs underwent apoptosis in livers 7 days after CCl_4_ cessation. We speculated the disappeared mGFP^+^ HSCs were due to apoptosis, and our immunohistochemical results approved that mGFP^+^ HSCs underwent apoptosis in livers recovered from CCl_4_-induced injury for 7 days (**Figure 7D**). Moreover, serial section immunohistochemistry showed 3.83% Desmin^+^ HSCs underwent apoptosis and 7.26% mGFP^+^ HSCs underwent apoptosis which was almost 2 times as high as the percentage of Desmin^+^ HSCs undergoing apoptosis (**Figure 7E**). These findings indicated that mGFP^+^ HSCs were more susceptible to apoptosis than Desmin^+^ HSCs.

**Figure 7.**
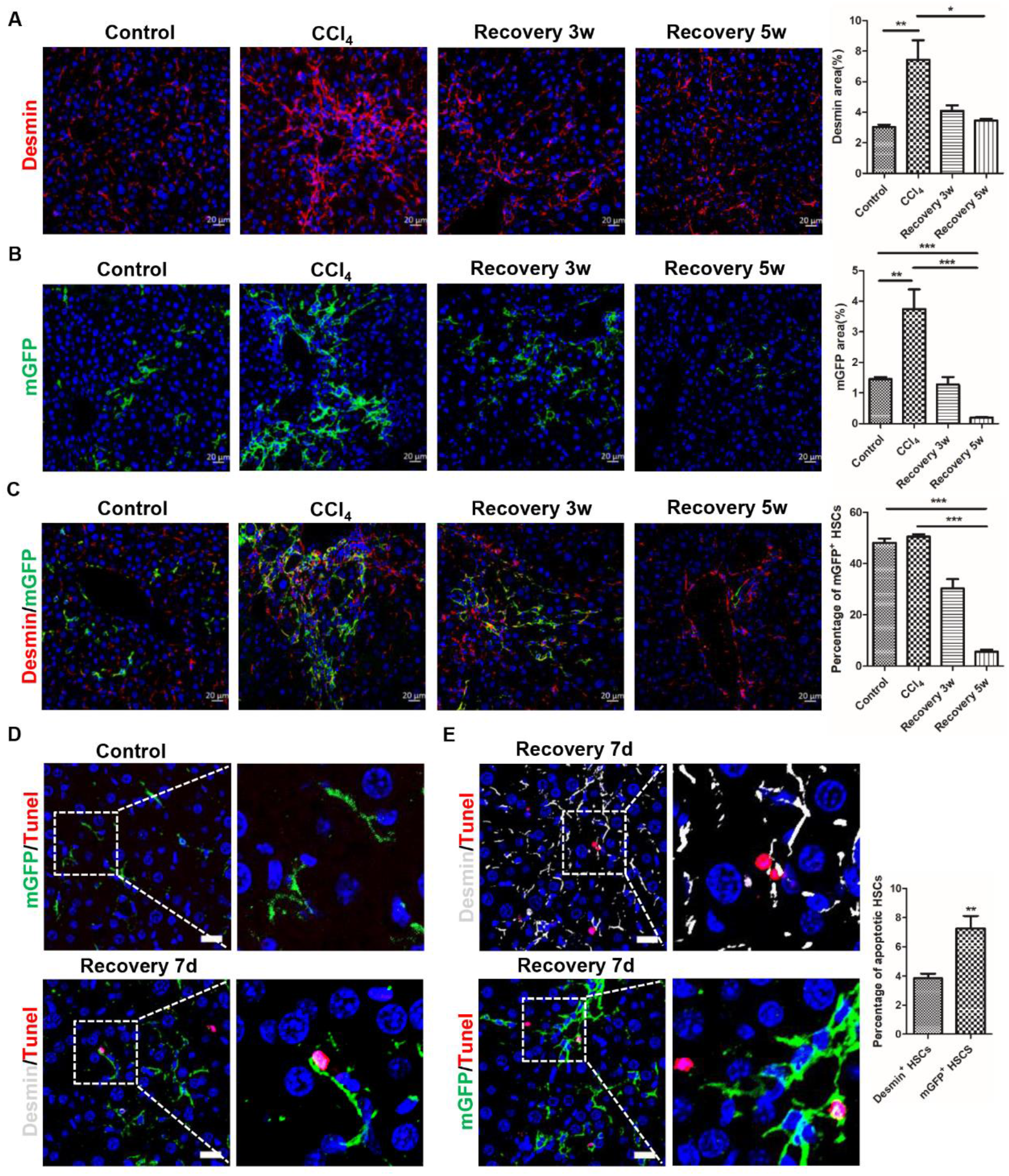
Reelin^+^ HSCs undergo apoptosis during recovery from CCl_4_-induced liver fibrosis. **A.** Immunostaining of Desmin in Reelin^CreERT2^; R26T/G^f^ mouse livers indicated that the number of Desmin^+^ HSCs recovered to normal level after CCl_4_ treatment was terminated for 5 weeks. **B.** Few mGFP^+^ HSCs existed after 5 weeks recovery from chronic CCl_4_ injury by analyzing immunohistochemistry of liver for mGFP. **C.** Immunohistochemistry for mGFP with Desmin observed that the percentage of mGFP^+^ HSCs accounted for Desmin^+^ HSCs was reduced significantly during recovery from CCl_4_-induced liver fibrosis. **D.** Apoptosed mGFP^+^ HSCs were observed by Tunel in Reelin^CreERT2^; R26T/G^f^ mouse livers recovery 7 days from CCl_4_-induced liver fibrosis. **E.** Immunohistochemistry for Tunel with mGFP or Desmin demonstrated that mGFP^+^ HSCs were more likely to undergone apoptosis than Desmin^+^ HSCs in serial sections. Data are reported as means ±SEM. *p < 0.05; **p< 0.01; ***p < 0.001. Scale bar represents 20 μm.

**Figure 8.**
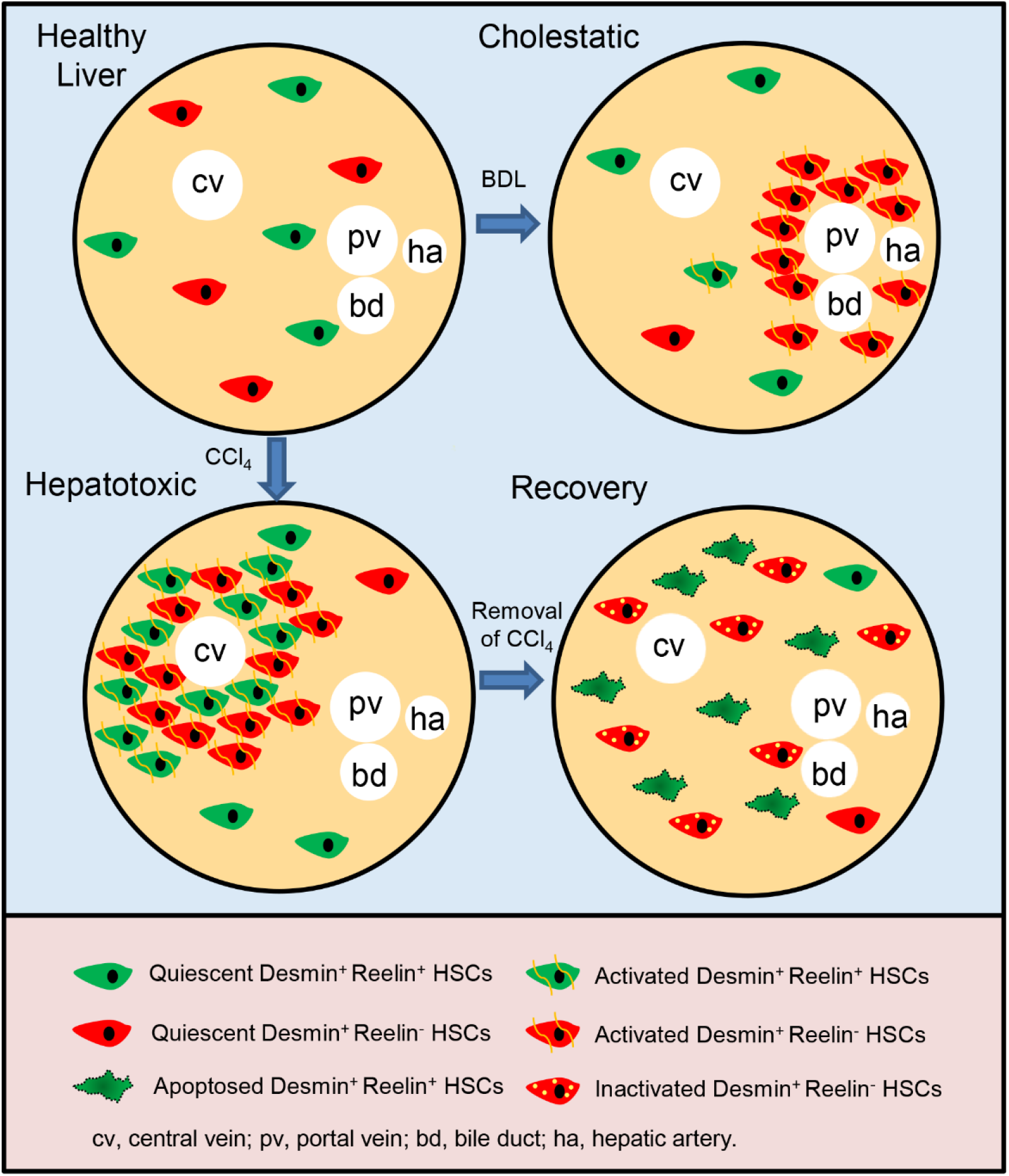
Graphical Abstract. - Cell lineage tracing reveals that about 50% HSCs are marked by ReelinCreERT2, and properties of these ReelinCreERT2-marked HSCs (Desmin^+^ Reelin^+^ HSCs) are different from Desmin^+^ Reelin^-^ HSCs.
- In cholestatic injury model, Desmin^+^ Reelin^-^ HSCs are accumulated around the portal triad with a significant activation and proliferation activity, but Desmin^+^ Reelin^+^ HSCs do not show proliferation or migration to the portal triad, and only a small part is activated.
- In hepatotoxic injury model, Desmin^+^ Reelin^+^ HSCs share similarities with Desmin^+^ Reelin^-^ HSCs in migration, proliferation and activation, nonetheless, still fewer Desmin^+^ Reelin^+^ HSCs are activated.
- In the regression of liver fibrosis, Desmin^+^ Reelin^+^ HSCs are apoptosed.

To ensure our findings that mGFP^+^ HSCs were a different subset compared to total HSCs, we chose Vimentin as another total HSC marker(17). Immunohistochemistry of Desmin and Vimentin in sham-operated and BDL-induced livers showed that Vimentin^+^ HSC’s characteristics were similar to Desmin^+^ HSC’s (**Supplemental Figure 3A**), and immunostaining of mGFP and Desmin or Vimentin showed that the percentage of mGFP accounted for Vimentin^+^ HSCs had no difference compared to Desmin+ HSCs (**Supplemental Figure 3B**). And the results of immunohistochemistry of mGFP and Desmin or Vimentin in normal and CCl_4_-treated livers were consistent with results in sham-operated and BDL-induced livers (**Supplemental Figure 3C and D**).

## Discussion

Reelin as a serine protease, play an important role in brain (42, 43). But which type of cells in livers expressing Reelin was controversial. Cell lineage tracing is a technique to track cell fate based on cre-lox system, revealing self-renewal, differentiation and migration of specific types of cells in development, disease and regeneration(23). Early studies applied cell lineage tracing to confirm a certain cell lineage is stem cells or investigate epithelial-to-mesenchymal transition (EMT) and MET(21). In our study, we pioneeringly changed the conventional usage of cell lineage tracing and applied this technique to demonstrate that Reelin was expressed in HSCs using Reelin^CreERT2^; R26T/G^f^ mouse model and these ReelinCreERT2-labled HSCs were a new subset, which displayed different characteristics compared to Desmin^+^ HSCs in vivo. We investigated properties of mGFP^+^ (Reelin^+^) HSCs and Desmin^+^ HSCs in activation, migration, and proliferation in BDL-induced and CCl_4_-induced fibrotic livers. Our results showed that in BDL-induced fibrotic livers, Desmin^+^ HSCs accumulated around the portal vein with significant proliferation and activation, whereas Reelin^+^ HSCs did not migrate either proliferate, and only a small part was activated compared to Desmin^+^ HSCs. HSC’s activation and proliferation usually are considered as a companying process(10), but our results indicate that the processes of activation and proliferation of HSCs are independent in some degree under certain conditions. And we observed that CCl_4_-Reelin^+^ HSCs exhibited similarity to CCl_4_-Desmin^+^ HSCs rather than BDL-Reelin^+^ HSCs. In CCl_4_-induced fibrotic livers, both of Desmin^+^ HSCs and Reelin^+^ HSCs accumulated around the central vein with remarkable proliferation activity, but the proliferative rate between Desmin^+^ and Reelin^+^ HSCs had no significant differences. The activation potential analysis of Desmin^+^ HSCs and Reelin^+^ HSCs showed Desmin^+^ HSCs and Reelin^+^ HSCs were both activated markedly, but fewer mGFP^+^ HSCs were activated compared to Desmin^+^ HSCs. Single-cell RNA sequencing is also a powerful method to identify a certain type of cell’s characteristics and has demonstrated distinct HSC clusters exist in normal or injured livers(9, 30). But physiological conditions are different from in vitro culture environments, for instance, cellular microenvironment, cytokines, cell-cell junction, et al, which might change the gene expression and chromatin state of HSCs(44, 45). So, single-cell RNA sequencing might get an incorrect result because of the difference of physiological conditions and in vitro culture environments. Compared to Single-cell RNA sequencing, our cell lineage tracing findings are closer to Reelin^+^ HSCs physiological properties in vivo.

To verify HSCs transforming into hepatocytes and cholangiocytes through MET, we traced Reelin^+^ HSC’s fate both in BDL-induced and CCl_4_-induced injured livers and we observed no Reelin^+^ HSCs expressed CK19 or HNF4α. In consideration of ReelinCreERT2 only marked about 50% HSCs, it is still possible that HSCs not marked by ReelinCreERT2 differentiate into cholangiocytes or hepatocytes through MET. Whether HSCs differentiating into cholangiocytes or hepatocytes through MET should be further investigated. Unexpectedly, in the regression of fibrosis, Desmin^+^ HSC’s number decreased to normal, but Reelin^+^ HSCs accounting for about 50% Desmin^+^ HSCs almost all disappeared because of apoptosis. Combining the early researches which reported that 50% of HSCs underwent inactivation during the liver injury recovery(28, 29), we speculated that Reelin^+^ HSCs underwent apoptosis and Reelin^-^ HSCs underwent inactivation in the regression of fibrosis.

We have not elucidated why there are marked differences in migration, activation and proliferation among BDL-Reelin^+^ HSCs, CCl_4_-Reelin^+^ HSCs, BDL-Desmin^+^ HSCs, and CCl_4_-Desmin^+^ HSCs. HSCs are main source of MFs independent of etiologies. In consideration of the heterogeneity of HSCs, properties of distinct clusters of HSCs are various in the same or different etiologies(30, 40). Portal fibroblasts (PFs) also are activated in BDL-induced liver injury, which are accumulated around the portal area and play a crucial part at the early stage of BDL-induced liver fibrosis(46, 47), and promote HSCs activation(18). Based on the findings that both of PFs and HSCs accumulating around the portal area(8, 47), and Reelin^+^ HSCs scattered throughout the parenchyma, we speculate PFs may have more effects on Desmin^+^ HSCs rather than Reelin^+^ HSCs though we did not investigate the interaction between PFs, Desmin^+^ HSCs, and Reelin^+^ HSCs. Additionally, the mechanisms of fibrogenesis are different in BDL and CCl_4_ induced liver injuries(35). BDL mainly results in obstruction of bile flow and increased biliary pressure which further gives rise to hyperplasia of biliary epithelial cells, causing cholestatic injury that progress to periportal fibrosis and cirrhosis(36). While CCl_4_ is hepatotoxic, leading to free radical reactions, lipid peroxidation, inflammatory response and necrosis of hepatocytes, and gives rise to an initial pericentral matrix deposition(38, 48). The differences in migration, activation and proliferation activities between Reelin^+^ HSCs and Desmin^+^ HSCs in BDL and CCl_4_ induced liver fibrosis maybe also caused by the model-specific influence.

In conclusion, we pioneeringly using cell lineage tracing have demonstrated that ReelinCreERT2-labled HSCs are a new cluster which displays different characteristics compared to Desmin^+^ HSCs in BDL-induced and CCl_4_-induced fibrotic livers. And our findings enlighten that treating liver fibrosis caused by different etiologies should suit the remedy to the case, for instance, focusing on Reelin^+^ HSCs may be a good therapy target in CCl_4_-induced liver fibrosis (hepatotoxic injury), however, controlling the Reelin^-^ HSCs may optimize the therapeutic effects in BDL-induced liver fibrosis (cholestatic injury).

## Materials and methods

### Mice

The animals in this study were against a C57BL6/J background. Rosa26mTmG reporter mice were obtained from Jackson Laboratory. ReelinCreERT2 mice were constructed by Biocytogen (Beijing, China). The P2A-iCreERT2 cassette was inserted after the stop codon TGA of Exon64 of Reelin and the knock-in mice were prepared based on the CRISPR/Cas9-based system developed by Biocytogen. ReelinCreERT2 mice were crossed with Rosa26mTmG reporter mice to generate Reelin^CreERT2^; Rosa26mTmG^flox^ (R26T/G^f^) mice used for subsequent experiments. ReelinCreERT2 genotype identification was performed by using forward primer 5’-CTCTGCTGCCTCCTGGCTTCT and reverse primer 5’-TCAATGGGCGGGGGTCGTT. Rosa26mTmG reporter mice genotype identification was conducted by using forward primer 5’-TATTCTGTCCCTAGGCGGTGAAGTCT and reverse primer 5’-CCTGTCCCTGAACATGTCCATCAG. All animal experiment procedures were in accordance with guidelines of Huazhong Agricultural University Guidelines for the Care and Use of Laboratory Animals.

### Fibrosis Induction and Tamoxifen Injection

Hepatic fibrosis was induced by intraperitoneal injections of carbontetrachloride (CCl_4_) (Aladdin, C112043, Shanghai, China) at the dose of 1ml/kg body weight, two times a week for 6-week (n=7), followed by 7-day (n=4), 3-week (n=6), or 5-week (n=6) recovery or induced by BDL for 2 weeks (n=5). CCl_4_ was dissolved in corn oil (Aladdin, C116023) at a ratio of 1:4. The number of mice treated with vehicle was 6 and the number of sham-operated mice was 3. ReelinCreERT2 activity was induced by intraperitoneal injections of tamoxifen (Sigma, T5648, Missouri, USA) at the dose of 100 mg/kg on daily basis for 3 days starting 7 days before the first CCl_4_ injection. Bromodeoxyuridine (BrdU) (Sigma, B5002) was injected at the dose of 50 mg/kg body weight every two hours for 4 times, the last injection was taken 24 hours before sacrifice.

### Tunel (terminal deoxynucleotidyl transferase dUTP nick end labeling)

We detected DNA fragmentation resulting from apoptotic signaling cascades with the In Situ Cell Death Detection Kit Fluorescein (Roche, 11684795910, Basel, Switzerland) according to manufacturer’s instructions. The presence of nicks in the DNA was identified by terminal deoxynucleotidyl transferase (TdT), an enzyme that catalyzed the addition of labeled dUTPs. The samples were digested with DNAse and used as staining positive control.

### Immunofluorescent Assay

Samples were fixed in 4% paraformaldehyde (PFA), embedded in paraffin, cut into 4 μm sections, dewaxed, hydrated, and subsequently incubated with antibodies. Fluorescence was bleached with 3% H2O2 in methanol for 15 minutes. For antigen retrieval, samples were heated in 10 mM sodium citrate buffer (pH 6.0) for 20 minutes. Sections were blocked with 10% goat serum for 30 minutes and incubated with primary antibodies, anti-GFP (Proteintech, 50430-AP, Wuhan, China), anti-GFP (Santa Cruz, sc-9996, Texas, USA), anti-desmin (Servicebio, GB12081, Wuhan, China), anti-GS(glutamine synthetase) (Santa Cruz, sc-74430), anti-CK19 (Servicebio, GB12197), and anti-HNF4α (Abcam, Ab41898, Cambridgeshire, UK), anti-Vimentin (Abcam, Ab92547), anti-Casp3 (Proteintech, 19677-1-AP), anti-BrdU (Servicebio, GB12051), anti-Ki67(Invitrogen, PA5-19462, Massachusetts, USA). Subsequently, sections were incubated with fluorophore-conjugated secondary antibodies (2.5 μg/ml, Invitrogen, A-11034, A-21424), nuclei co-staining with 4, 6-diamidino-2-phenylindole (DAPI) (Abcam, ab104139). Images were acquired with a laser scanning confocal microscope (Carl Zeiss Microscopy, LSM710, Jena, Germany), and were analyzed by Zen software with fixed parameters.

### Histology and immunohistochemistry

Liver tissues were immobilized with 4% PFA, dehydrated, embedded in paraffin, sectioned at 4 μm, and processed for Sirius red staining and immunohistochemistry. Immunohistochemistry was performed with antibodies, anti-GFP (Santa Cruz, sc-9996) and anti-Reelin (NOVUS, NB600-1081, Minneapolis, USA). Subsequently sections were incubated with diaminobenzidine (Gene Tech Company Limited, GK347011, Shanghai, China) and counterstaining with hematoxylin (Servicebio, GB1004). All steps of immunohistochemistry are according to manufacturer’s instructions.

### Software–Intensity measurement

Image Pro Plus (Image Pro Plus v.7: Media Cybernetics; Bethesda, MD), as an analysis program, was used to analyze and quantify data from photomicrographs. In this study, the analyses were performed as follows: Integrated Optical Density (IOD) Image Pro Plus was used to quantify the intensity of probes binding to the structures. We used the confocal series to calculate the total binding intensity of the probes (IOD-intensity value).

### Statistical analysis

Statistical analyses were performed using the GraphPad Prism 6(GraphPad). Data are expressed as means ± SEM. Comparisons between two groups were performed using the two-tailed Student’s t-test. Comparisons between multiple groups were performed using ordinary one-way ANOVA with the Dunnett’s multiple comparison test. Statistical significance was presented at the level of *p < 0.05, **p < 0.01, ***p < 0.001.

## Acknowledgements

This work was supported by National Key R&D Plan no. 2017YFA0103202 and no. 2017YFA0103200, National Natural Science Foundation of China 32071143, the Fundamental Research Funds for the Central Universities (2662019YJ008), and Huazhong Agricultural University Startup funds. We are thankful to Nannan Niu, and Bingjie Li (Huazhong Agricultural University) for critical reading of the manuscript.

## Author contributions

N. C. and L. Z. conceived and designed the study; N. C. provided the experimental data; N.C. and S.L. performed the experiments; D.Q., D.G., Y.C., C.H., and S.Z. provided assistance in animal experiments; N.C. and L. Z. discussed and drafted the manuscript; L. W. and X. C reviewed the manuscript; L.Z and N.C. organized the data and wrote the manuscript.

## Ethic statement

All animal experiments conducted were compliant with Huazhong Agricultural University Guidelines for the Care and Use of Laboratory Animals

## Data availability

The data that support the findings of this study are available from the corresponding author upon reasonable request.

## Supplemental Figures

**Supplemental Figure 1.**
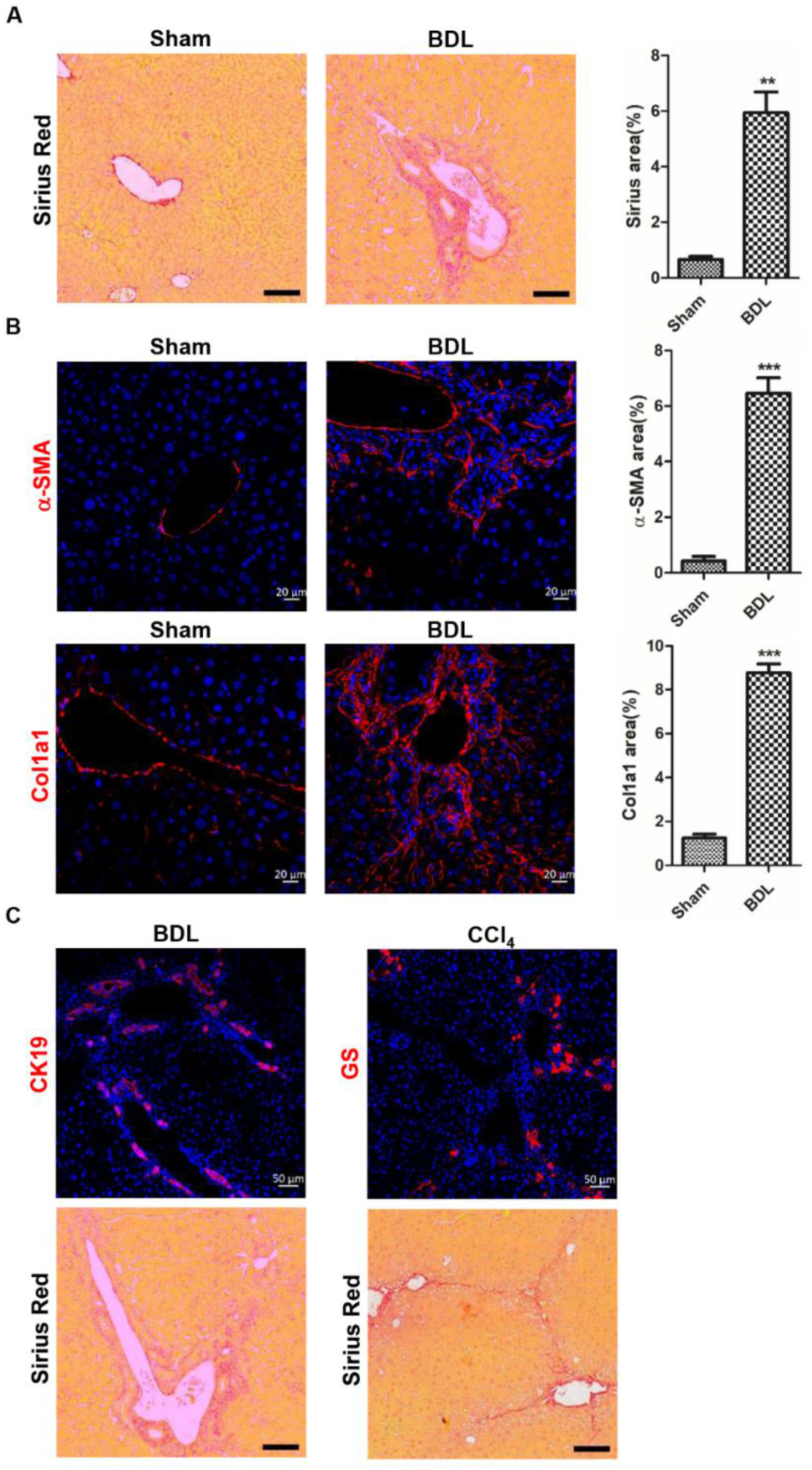
BDL induces severe biliary fibrosis. **A.** Sirius red staining displayed that severe biliary fibrosis was induced by BDL for 2 weeks in Reelin^CreERT2^; R26T/G^f^ mice**. B.** α-SMA and Col1a1 expression was significantly increased in BDL-operated Reelin^CreERT2^; R26T/G^f^ mice**. C.** Immunohistochemistry for CK19, GS and Sirius red indicated that BDL caused biliary duct hyperplasia and biliary fibrosis, whereas CCl_4_ led to pericentral cell injury and fibrosis and forming fibrous septum. Data are reported as means ± SEM. *p < 0.05; **p< 0.01; ***p < 0.001. Scale bar in A represents 100 μm. Scale bar in B represents 20 μm. Scale bar in C immunohistochemistry pictures represents 50 μm and in Sirius red staining represents 100 μm.

**Supplemental Figure 2.**
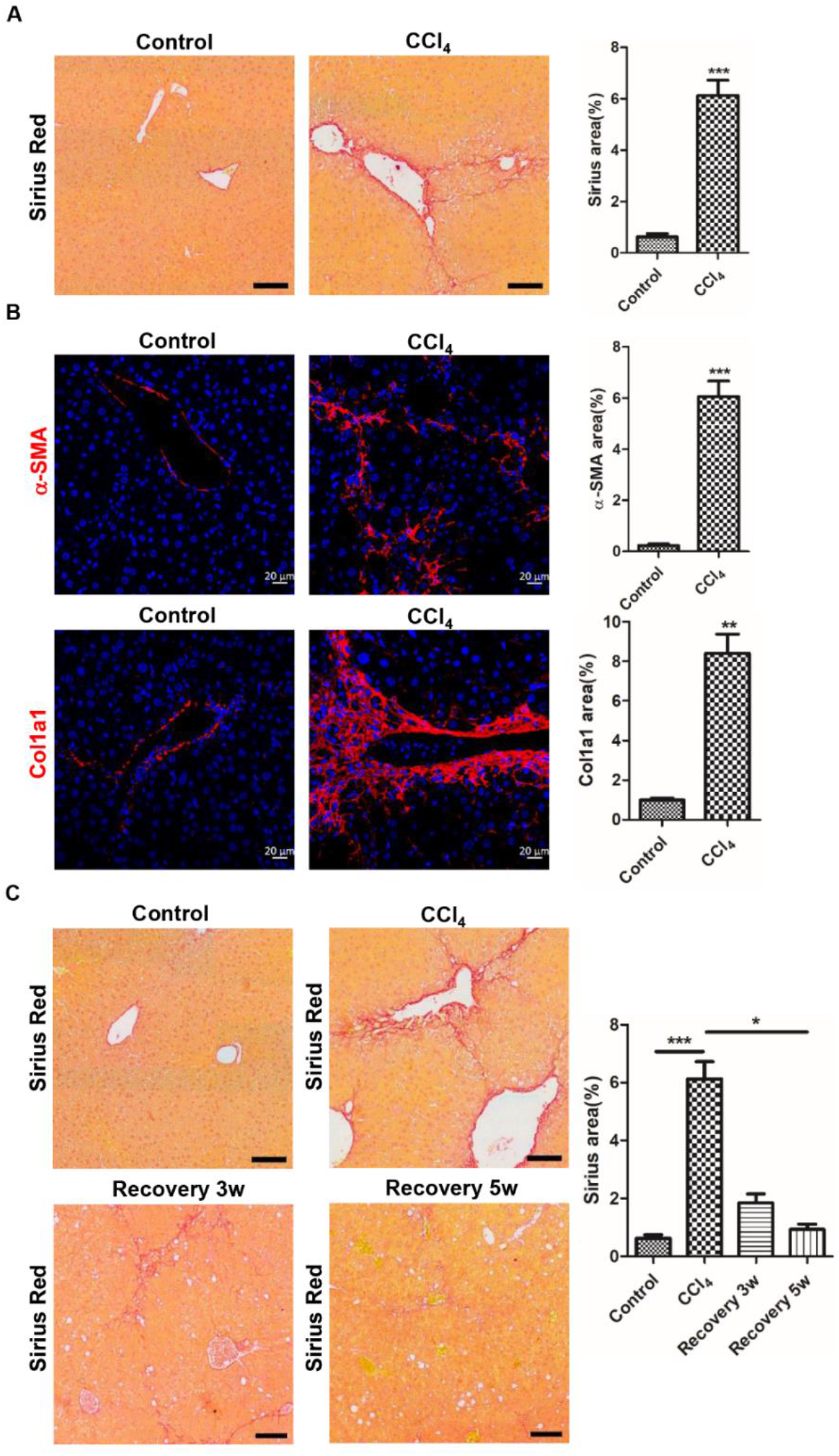
CCl_4_ induces severe pericentral fibrosis. **A.** Severe pericentral fibrosis was induced by CCl_4_ for 6 weeks in Reelin^CreERT2^; R26T/G^f^ mice observed by sirius red staining**. B.** α-SMA and Col1a1 expression were significantly increased in CCl_4_-treated Reelin^CreERT2^; R26T/G^f^ mice for 6 weeks**. C.** Sirius red staining indicated that after 5 weeks recovery from CCl_4_, fibrotic livers recovered to normal state. Data are reported as means ±SEM. *p < 0.05; **p< 0.01; ***p < 0.001. Scale bar in A and C represents 100 μm. Scale bar in B represents 20 μm.

**Supplemental Figure 3.**
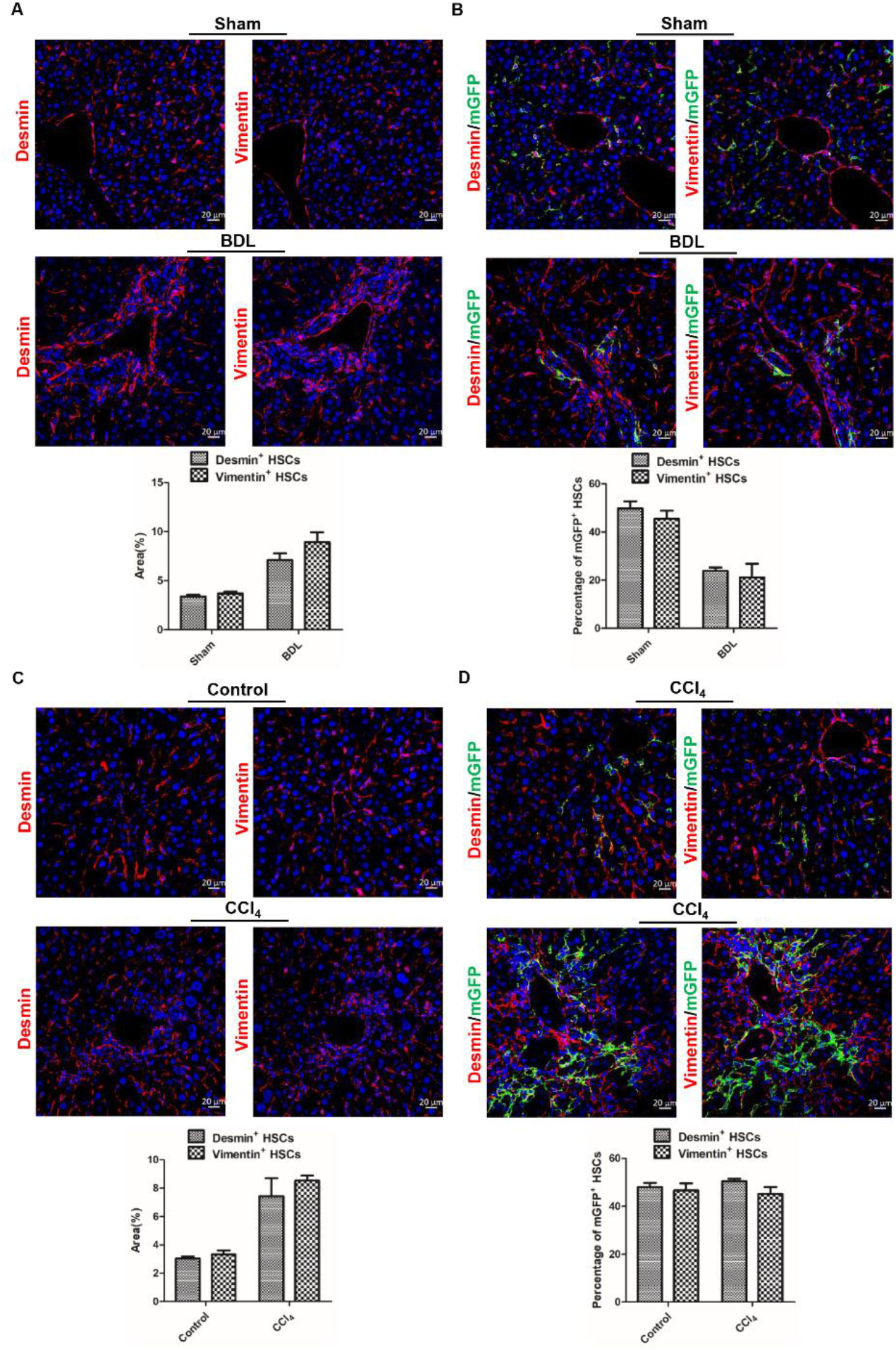
The characteristics of Vimentin^+^ HSCs are consistent with Desmin^+^ HSCs in BDL-induced and CCl_4_-induced liver fibrosis. **A.** Serial sections immunostaining of Vimentin and Desmin indicated that the number and distribution of Vimentin^+^ HSCs were consistent with Desmin^+^ HSCs in sham-operated and BDL-operated Reelin^CreERT2^; R26T/G^f^ mouse livers. **B.** Serial sections immunostaining of mGFP and Desmin or Vimentin determined that the percentage of mGFP^+^ HSCs accounted for Desmin^+^ HSCs was similar to accounted for Vimentin^+^ HSCs in sham-operated and BDL-operated Reelin^CreERT2^; R26T/G^f^ mouse livers. **C.** Serial sections immunostaining of Vimentin and Desmin indicated that the number and distribution of Vimentin^+^ HSCs were consistent with Desmin^+^ HSCs in normal and CCl_4_-treated Reelin^CreERT2^; R26T/G^f^ mouse livers. **D.** Serial sections immunostaining of mGFP and Desmin or Vimentin indicated that the percentage of mGFP^+^ HSCs accounted for Desmin^+^ HSCs was similar to accounted for Vimentin^+^ HSCs in in normal and CCl_4_-treated Reelin^CreERT2^; R26T/G^f^ mouse livers. Data are reported as means ± SEM. *p < 0.05; **p< 0.01; ***p < 0.001. Scale bar represents 20 μm.

